# Flow cytometric analysis and purification of airway epithelial cell subsets

**DOI:** 10.1101/2020.04.20.051383

**Authors:** Luke R. Bonser, Kyung Duk Koh, Kristina Johansson, Semil P. Choksi, Dan Cheng, Leqian Liu, Dingyuan I. Sun, Lorna T. Zlock, Walter L. Eckalbar, Walter E. Finkbeiner, David J. Erle

## Abstract

The human airway epithelium is essential in homeostasis, and epithelial dysfunction contributes to chronic airway disease. Development of flow cytometric methods to characterize subsets of airway epithelial cells will enable further dissection of airway epithelial biology. Leveraging single cell RNA-sequencing (scRNA-seq) data in combination with known cell type-specific markers, we developed panels of antibodies to characterize and isolate the major airway epithelial subsets (basal, ciliated, and secretory cells) from human bronchial epithelial cell cultures. We also identified molecularly distinct subpopulations of secretory cells and demonstrated cell subset-specific expression of low abundance transcripts and micro-RNAs that are challenging to analyze with current scRNA-seq methods. These new tools will be valuable for analyzing and separating airway epithelial subsets and interrogating airway epithelial biology.

## Introduction

Flow cytometry is a commonly used research and diagnostic tool that uses fluorophore-conjugated antibodies as probes to identify, characterize, and/or isolate cell populations (1). The immunology community has developed panels of antibodies useful for detailed immunophenotyping of immune cells derived from many organs including the lung. A recent American Thoracic Society working group report (2) heralded the importance of flow cytometry in pulmonary research, but noted that “the development of appropriate markers for nonimmunologic cells is less mature than other pulmonary cell types,” including epithelial cells.

The airway epithelium defends against inhaled environmental challenges including pollutants, pathogens, and allergens (3), and epithelial dysfunction is central to the pathogenesis of major lung diseases including asthma, cystic fibrosis, chronic obstructive pulmonary disease, and primary ciliary dyskinesia (4). Proximal airway epithelial subsets including basal cells, ciliated cells, secretory cells including club and goblet cells, intermediate cells, brush cells, and pulmonary neuroendocrine cells have been defined morphologically and by their anatomical location within the tissue by histology and electron microscopy (5, 6). Single cell RNA-sequencing (scRNA-seq) is further advancing our understanding of airway epithelial heterogeneity. Recent studies of human and murine airways have confirmed the presence of previously defined major airway epithelial subsets, identified molecularly distinct subpopulations of these cells, and uncovered previously unrecognized cell types including ionocytes (7–11).

Pan-epithelial antibodies (EpCAM/pan-cytokeratin (12)) and limited sets of cell type-specific antibodies (e.g., acetylated alpha-tubulin (TUBA) for ciliated cells (13), nerve growth factor receptor (NGFR) and integrin subunit alpha 6 (ITGA6) (14) for basal cells, and mucin-5AC (MUC5AC) for goblet cells (15)) have been used individually for flow cytometry. We sought to develop a larger panel of antibodies for simultaneous analysis of the major subsets of airway epithelial cells and provide a new tool for further understanding airway epithelial physiology and pathology. To this end, we used known markers described elsewhere in the literature, augmented these by leveraging information from scRNA-seq datasets, and developed panels of antibodies for flow cytometry to identify, characterize, and isolate the major human airway epithelial subsets.

## Methods

Additional details are provided in the online data supplement. A detailed protocol is provided in the online supplementary document.

### Primary human bronchial epithelial cell (HBEC) culture

HBECs isolated from lungs not used for transplantation were cultured at air-liquid interface (ALI) as previously described (16, 17). We harvested cells 23 days after establishment of ALI. Some cultures were stimulated with IL-13 10 ng/ml for the final seven days of culture to induce goblet cell production as indicated (18). The UCSF Committee on Human Research approved the use of HBECs for these studies.

### Flow cytometry analysis

We trypsinized HBECs to generate single cell suspensions and fixed cells in 0.5% (v/v) paraformaldehyde; if not stained immediately, cells were frozen at −80 °C. Cells were blocked, stained with the analytical panel (Table E1), and analyzed by flow cytometry.

### Flow cytometry cell sorting

Prior to trypsinization, cells were incubated in culture media containing SiR-Tubulin (Spirochrome/Cytoskeleton Inc., Denver, CO). Single cell suspensions were generated as above, and singlets stained with fixable viability dye eFluor450 (ThermoFisher Scientific, Fremont, CA, USA) to discriminate live cells. Cells were subsequently stained with a sorting panel comprising cell surface markers only (Table E1) and sorted by flow cytometry.

### Gene expression analysis

We isolated total RNA from sorted cells, performed reverse transcription, and analyzed the cDNA by quantitative real-time PCR (qRT-PCR) to quantify specific mRNAs and microRNAs (miRNAs). Table E2 lists qRT-PCR primer sequences.

## Results

### Identification of airway epithelial subset markers

To identify a panel of candidate cell-subset specific markers, we combined markers previously used for flow cytometry (TUBA, ITGA6, NGFR, and ITGA6) with transcripts identified in several recent human scRNA-seq datasets (cadherin-related family member 3 (CDHR3) and CEA cell adhesion molecule (CEACAM) 5) (7, 9–11). We also analyzed HBECs differentiated at air-liquid interface (16, 17) using the Drop-seq scRNA-seq platform (19); IL-13-stimulated cultures were included as IL-13 induces goblet cell production (18) (Figure E1B). We examined our dataset for cell-type–specific transcripts (Figure E1B), defined as genes more highly expressed in one cell type than the others (FDR < 0.05) and identified two additional markers (CEACAM6 and tetraspanin-8 (TSPAN8)). We combined all markers into a prospective flow panel (Table 1).

**Table 1.** Panel of cell-subset-specific markers

| Name | Antigen | Cell subset | Description | References |
| --- | --- | --- | --- | --- |
| Acetylated alpha-tubulin | TUBA | Ciliated | Recognizes axonemal alpha-tubulin acetylated on the epsilon-amino group of lysine(s), a hallmark of stable microtubules, which are enriched in motile cilia | (7, 9, 10, 21, 29) |
| Cadherin-related family member 3 | CDHR3 | Ciliated | Calcium-dependent cell adhesion protein associated with ciliogenesis and asthma; receptor for Rhinovirus C | (9, 10, 13, 30, 3) |
| integrin subunit alpha 6 | CD49f/<br>ITGA6 | Basal | Integral membrane protein of the integrin alpha chain family; function uncharacterized in the airway | (10, 14) |
| Nerve growth factor receptor | CD271/<br>NGFR | Basal | Cell surface receptor localized to basal cells of unknown function in the airway | (7, 14) |
| CEA cell adhesion molecule 5 | CD66c/<br>CEACAM5 | Secretory | Cell surface glycoprotein upregulated after smoking cessation and linked to lung squamous cell carcinoma | (9, 10, 32, 33) |
| CEA cell adhesion molecule 6 | CD66e/<br>CEACAM6 | Secretory | Cell surface glycoprotein associated with severe asthma and upregulated after smoking cessation | (7, 9, 10, 32, 34) |
| Tetraspanin 8 | TSPAN8 | Goblet | Cell surface glycoprotein of unknown function in the airway | (9, 10) |
| Mucin 5AC | MUC5AC | Goblet | Secreted, gel-forming glycoprotein associated with mucus dysfunction in chronic lung disease | (4, 9, 10) |

To test whether these putative cell subset markers were suitable for flow cytometry, we stained unstimulated and IL-13–stimulated HBECs from 5 donors with antibodies against these markers individually and performed flow cytometry. Each antibody stained a subset of HBECs from both unstimulated and IL-13–stimulated cell cultures, except for the goblet cell markers TSPAN8 and MUC5AC, which stained a subset of cells from IL-13–stimulated cultures, but few if any cells from unstimulated cultures (Figure 1A). We observed significant increases in cells staining for TSPAN8 (*P* = 0.022), and MUC5AC (*P* = 0.0016) following IL-13-stimulation; however, we did not observe statistically significant effects of IL-13-stimulation on the proportion of cells stained for the other markers (Figure 1B).

**Figure 1:**
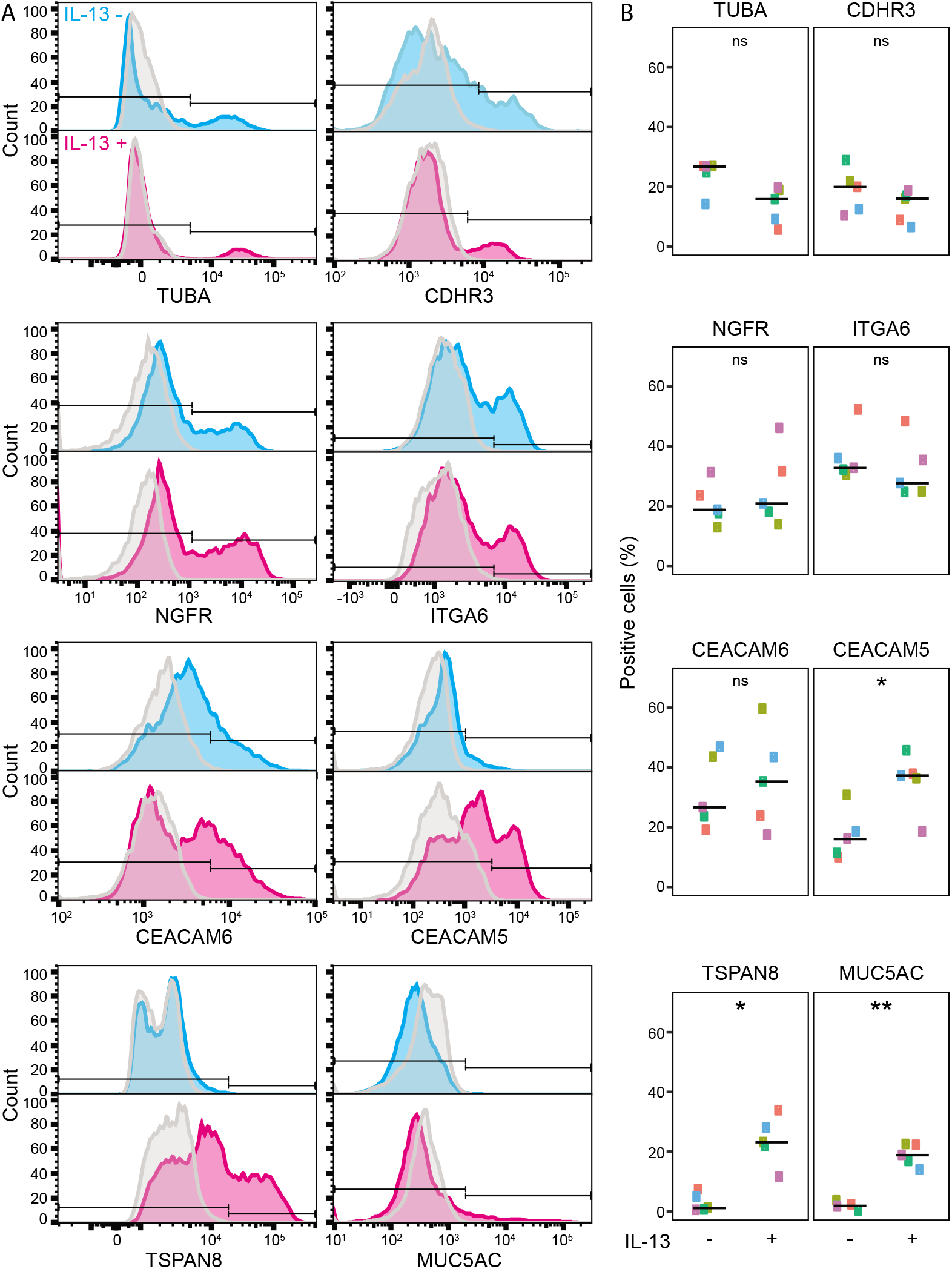
Identification of cell-specific markers for flow cytometry. (*A*) Human bronchial epithelial cells (HBECs) grown in the absence (cyan histogram) or presence of interleukin-13 (IL-13; magenta histogram) were stained using antibodies against cell subset specific markers. Positive staining was determined by comparing the same sample stained with an appropriate isotype control (grey histogram). (*B*) Comparison of positive staining for cell-subset–specific markers from unstimulated and IL-13–stimulated HBECs (n=5; each donor is represented by a different color). Significance was evaluated using a *t*-test: ns, nonsignificant; *, *P* < 0.05; and **, *P* < 0.01.

### Characterization of airway epithelial cells subsets using an analytical antibody panel

To define the relationships between cell subsets defined by these eight individual markers, we performed multicolor flow cytometry with an analytical panel comprising all eight antibodies (Table E1) on unstimulated and IL-13–stimulated HBECs from three donors. This produced a 10-dimensional dataset (one fluorescence intensity for each of the eight antibodies, forward scatter (FSC), and side scatter (SSC)). We analyzed this dataset with t-Distributed Stochastic Neighbor Embedding (tSNE; Figure 2A-J). The distribution of marker staining indicated that cell type heterogeneity was the major driver of staining patterns in this data set, although effects of IL-13-stimulation (Figure 2K) and inter-donor variation (Figure 2L) also contributed to the staining patterns. We next examined each staining parameter in relation to the tSNE plot. TUBA staining (Figure 2A) was confined to a distinct cluster of cells that were predominantly CDHR3^+^ (Figure 2B); cells bearing these two ciliated cell markers had little if any basal or secretory cell marker staining. Another cluster comprised cells stained for the basal cell markers NGFR (Figure 2C) and ITGA6 (Figure 2D), but with little if any secretory or ciliated cell marker staining, except for a small subset that stained for CDHR3. Cells that stained for neither ciliated nor basal cell markers generally stained with antibodies against one or both of the secretory cell markers CEACAM5 (Figure 2E) and CEACAM6 (Figure 2F). CEACAM5 staining was more pronounced in IL-13–stimulated cells. TSPAN8 and MUC5AC were detected only after IL-13 stimulation. TSPAN8 staining was limited to a subset of CEACAM5^+^ cells (Figure 2G); MUC5AC staining was confined to a subset of the TSPAN8^+^ population (Figure 2H). Ciliated cells (TUBA^+^ and CDHR3^+^) and some secretory cells tended to have higher forward scatter (Figure 2I), an indicator of cell size, and basal cells (ITGA6^+^ and frequently NGFR^+^) tended to have lower side scatter (Figure 2J).

**Figure 2:**
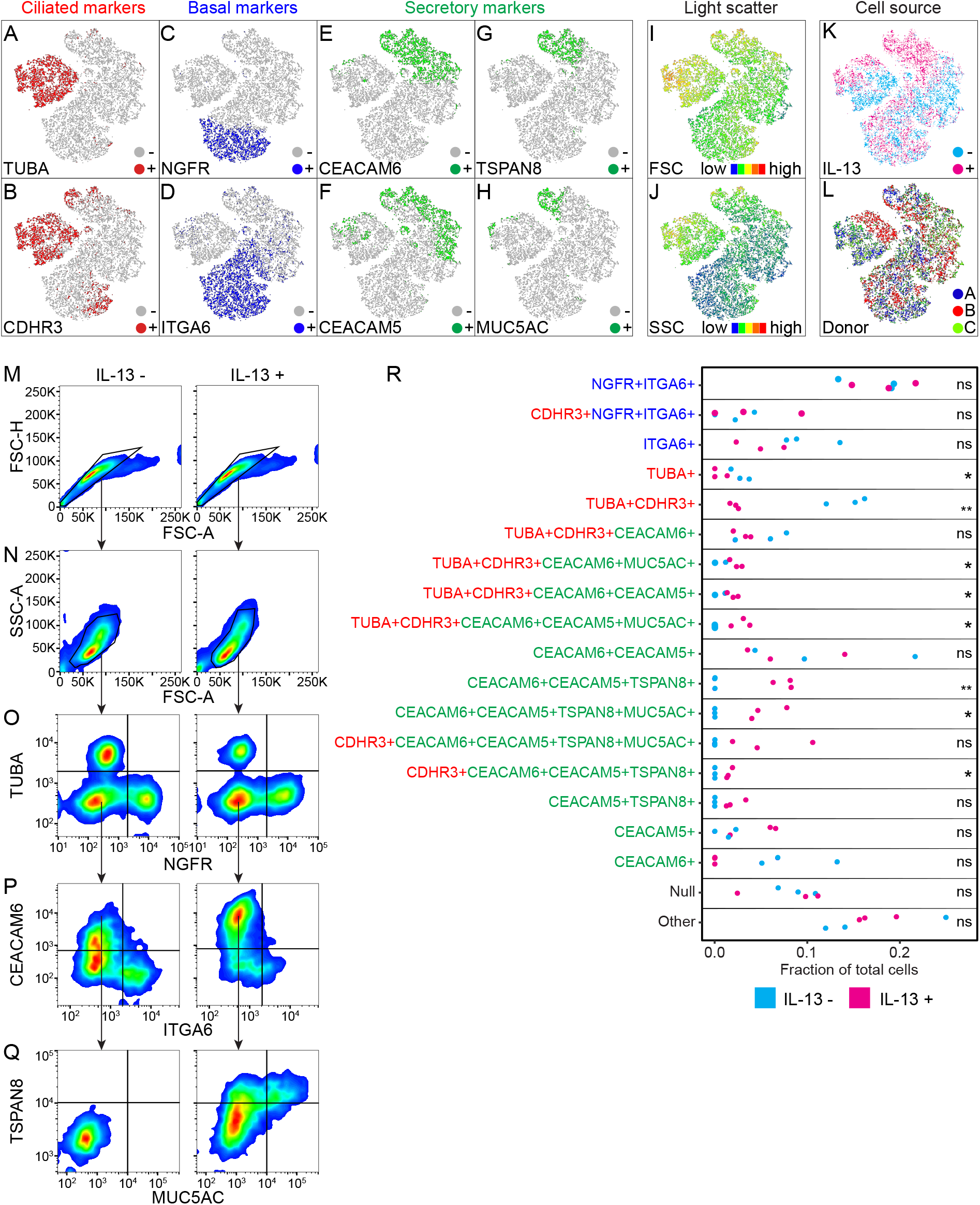
Characterization of airway epithelial cells subsets and IL-13 stimulation using an analytical flow cytometry panel. HBECs (n=3) were cultured with or without IL-13 and then processed for multicolor flow cytometry. (*A-L*) Results from 3,000 cells from each donor were combined and analyzed by t-Distributed Stochastic Neighbor Embedding (tSNE). tSNE plots are colored to show cells that stained for the ciliated cell markers (red) TUBA (*A*) and CDHR3 (*B*), the basal cell markers (blue) NGFR (*C*) and ITGA6 (*D*), and the secretory cell markers (green) CEACAM6 (*E*), CEACAM5 (*F*), TSPAN8 (*G*), and MUC5AC (*H*). Forward scatter FSC (*J*) and side scatter SSC (*K*) are represented as a continuum from low (blue) to high (red). Cells came from unstimulated (-) or IL-13-stimulated cultures (*K*) from three donors (*L*). (*M-Q*) Gating strategy to identify airway epithelial subsets from unstimulated (IL-13-) and IL-13–stimulated (IL-13+) HBECs. Doublets (*M*) and debris (*N*) were removed, and resulting singlets were gated on NGFR and TUBA (*O*). Subsequently, TUBA^-^NGFR^-^ singlets were gated on ITGA6 and CEACAM6 (*P*). ITGA6^-^CEACAM6^+^ cells were then gated on MUC5AC and TSPAN8 (*Q*). (*R*) Comparison of cell subsets (>1%) from unstimulated (IL-13-; cyan) and IL-13–stimulated (IL-13+; magenta) HBECs (n=3; each donor is represented by a different point). Ciliated cell markers are colored blue, basal cell markers red, and secretory cell markers green. “Null” indicates cells that did not stain with any of the eight markers and “Other” indicates totals for all subsets representing <1% of total cells. Significance was evaluated using Student’s *t*-test: ns, not significant; *, *P* < 0.05; **, *P* < 0.01.

To explore cellular heterogeneity, we examined the combination of the eight analytical panel markers on individual cells. Figure 2M-Q and Figure E2 illustrate the use of a sequential gating strategy to identify subpopulations of cells. Cells expressing the ciliated cell marker TUBA were clearly distinct from those expressing the basal cell marker NGFR, and cells lacking both TUBA and NGFR included subsets of cells expressing various combinations of the secretory cell markers CEACAM6, TSPAN8, and MUC5AC that were markedly altered by IL-13 stimulation.

We considered all 2^8^ (256) possible combinations of staining that could be seen using the panel of 8 markers and focused our attention on all subsets containing at least 1% of total cells (Figure 2R and Table E3). In unstimulated HBECs, 12 such subsets were observed. After IL-13 stimulation, we identified 17 subsets, 11 of which were present in unstimulated culture conditions. Subsets expressing basal cell markers (NGFR^+^ITGA6^+^, CDHR3^+^NGFR^+^ITGA6^+^, and ITGA6^+^ only) were identified in both culture conditions and their proportions were not significantly changed by IL-13. TUBA^+^CDHR3^+^ cells and TUBA^+^ only cells were also present in both unstimulated and IL-13-stimulated cultures, although the TUBA^+^ only subset was significantly reduced following IL-13-stimulation. Four TUBA^+^CDHR3^+^ cell subsets that co-stained with combinations of the secretory cell markers CEACAM6, CEACAM5, and MUC5AC were also observed; three of these subsets were significantly increased by IL-13. Cells expressing both ciliated and secretory cell markers have been also identified in scRNA-seq experiments, with pseudotime analyses suggesting that these cells may represent a transition from secretory to ciliated cells (11). Of cells bearing secretory cell markers, CEACAM6^+^CEACAM5^+^ cells were present in similar proportions in unstimulated and IL-13-stimulated conditions, as were CEACAM5^+^ only cells. CEACAM6^+^ only cells were unique to unstimulated cultures. IL-13 stimulation led to the emergence of cells expressing both

CEACAM5 and TSPAN8. These cells could be further subdivided into five subsets based on expression of other markers. Three of the five CEACAM5^+^TSPAN8^+^ subsets also stained with MUC5AC, as would be expected for goblet cells. Whether the CEACAM5^+^TSPAN8^+^MUC5AC^-^ populations represent goblet cell precursors, goblet cells that have secreted MUC5AC, or novel IL-13–induced secretory cell subsets requires further investigation. Our findings are consistent with bronchial tissue staining that demonstrated colocalization of CEACAM5 and MUC5AC in never and current smokers (10). We also identified a subset that stained with none of the markers (Null) and was present under both culture conditions. Collectively, these data suggest IL-13 alters epithelial cell diversity, shifting populations towards cell subsets exhibiting secretory and goblet cell markers.

### A modified flow cytometry panel is suitable for live cell sorting

Flow cytometry is also useful for live cell sorting. Our analytical panel precludes this since it requires fixation and permeabilization for staining with antibodies against the intracellular antigens MUC5AC and TUBA. We therefore designed a sorting panel that omitted the MUC5AC antibody and replaced the TUBA antibody with SiR-tubulin, a membrane-permeable live cell dye that stains microtubules (20), which have a major structural role in cilia (21). To ensure that we only sorted live cells, we also included a viability dye. The sorting panel is detailed in Table E1.

To evaluate this panel, we performed flow cytometric cell sorting on unfixed HBECs from 3 donors (Figure 3A-E). After excluding dead cells that failed to exclude the viability dye, we gated on SiR-tubulin and NGFR and identified the SiR-tubulin^+^NGFR^-^ singlets. Sorted SiR-tubulin^+^NGFR^-^ cells were subsequently stained for TUBA and examined by microscopy; 58/60 cells examined possessed cilia and stained with TUBA, confirming that these were ciliated cells (Figure 3F and G). We gated the remaining singlets on CEACAM6 and NGFR staining to discriminate between secretory and basal cell subpopulations.

**Figure 3:**
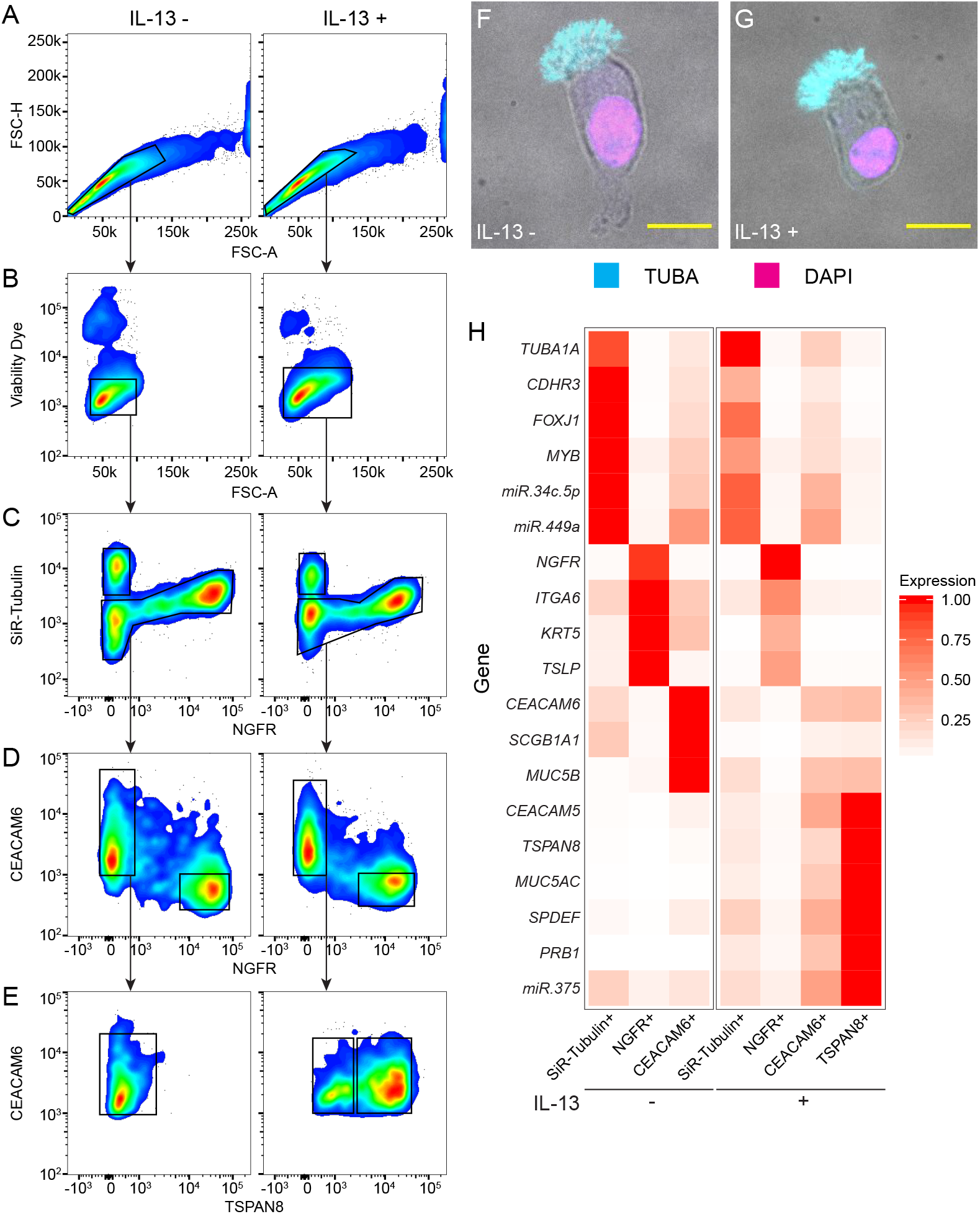
Flow cytometric sorting of HBEC subsets. Prior to processing for flow cytometry, HBECs cultured in the absence or presence of IL-13 were stained with SiR-tubulin. SiR-tubulin– stained cells were trypsinized, stained with a cell viability dye, and sorted by FACS. (*A-E*) Gating strategy for flow cytometric cell sorting of unstimulated (IL-13-) and IL-13–stimulated (IL-13+) HBECs. After selection of singlets (*A*) and live cells (*B*), cells were gated on SiR-tubulin and NGFR staining (*C*). SiR-tubulin^-^ cells were gated on NGFR and CEACAM6 (*D*). NGFR^-^ CEACAM6^+^ cells were gated on MUC5AC and TSPAN8 (*E*). (*F, G*) To validate SiR-tubulin staining, SiR-tubulin^+^ cells were fixed in paraformaldehyde, immobilized to slides by cytospin, stained with TUBA (cyan), and counterstained with DAPI (blue). Images show cells from unstimulated (IL-13-, *F*) and IL-13–stimulated (IL-13+, *G*) cultures with TUBA-stained cilia (found in 58/60 cells examined). Scale bar = 10 μM. (*H*) Quantitative real-time PCR analysis of sorted cell subpopulations. Mean expression values calculated from triplicate experiments with different donors were normalized to the maximum expression of the gene in any cell type (0-1: white-red).

We sorted these three cell subsets from HBECs, isolated total RNA, and performed qRT-PCR for cell type-specific markers (Figure 3H). As expected, the ciliated cell transcripts forkhead box J1 (*FOXJ1*), tubulin alpha 1A class 1 (*TUBA1A*), and *CDHR3* were enriched in the SiR-tubulin^+^ ciliated cell subset. The basal cell transcripts cytokeratin 5 (*KRT5*) and *NGFR* were enriched in the NGFR^+^ basal cell subset; this subset also expressed *ITGA6*. The secretory cell transcripts, secretoglobulin family 1A member 1 (*SCGB1A,* also known as club-cell specific protein, *CCSP*) and mucin-5B (*MUC5B*), along with *CEACAM6*, were enriched in the CEACAM6^+^ secretory cell subset, as expected. The goblet cell transcripts SAM pointed domain containing ETS transcription factor (*SPDEF*), *MUC5AC*, and *TSPAN8* were upregulated in CEACAM6^+^TSPAN8^+^ cells from IL-13– stimulated HBEC cultures. Collectively, this analysis demonstrated that the combination of cell surface marker and SiR-tubulin staining was sufficient for discriminating the major airway epithelial cell populations.

Analyzing sorted cells may improve the ability to detect low abundance transcripts that are difficult to quantify using available scRNA-seq approaches. We identified several transcripts detected in a bulk RNA-seq dataset (22) but absent from our scRNA-seq dataset, and performed qRT-PCR to determine their cell-subset–specific expression. For example, the transcription factor *SPDEF* is critical for IL-13–induced goblet cell differentiation of HBECs (15) but was not detected in our scRNA-seq dataset. Using our sorting panel, we found that *SPDEF* was selectively expressed in CEACAM6^+^ secretory cells, particularly CEACAM6^+^TSPAN8^+^ cells (Figure 3D). The alarmin thymic stromal lymphopoietin (*TSLP*) was almost exclusively expressed in the NGFR^+^ basal cell subset. The ciliated cell transcription factor MYB protooncogene, transcription factor (*MYB*) (23) was expressed primarily in SiR-tubulin^+^ ciliated cells and was downregulated by IL-13. Expression of the secreted protein, proline-rich protein BstNI subfamily 1 (*PRB1*), which we identified as an IL-13-induced gene by bulk RNA-seq but did not detect using Drop-seq, was enriched in the CEACAM6^+^TSPAN8^+^ secretory cell subset.

Analyzing sorted cells also allows for analysis of small RNAs, such as miRNAs, that are not assessed using Drop-seq and related scRNA-seq approaches. miR-34/449 family miRNAs are required for motile ciliogenesis (24). We quantified miR-34c-5p and miR-449a and confirmed enrichment in SiR-tubulin^+^ ciliated cells. We found that miR-375, which is involved in goblet cell differentiation in the colon (25), was IL-13–inducible, restricted to secretory cells, and enriched in the TSPAN8^+^ secretory cell subset. These data demonstrate that the sorting strategy we developed is useful for isolating and characterizing subpopulations of epithelial cells with distinct transcriptional and miRNA profiles.

## Discussion

Our study outlines an analytical flow panel and gating strategy for the characterization and enumeration of subsets of airway epithelial cells from HBECs. We also demonstrate a scheme for isolating common airway epithelial subsets from HBECs using a sorting flow panel. Both panels identified the major airway epithelial subsets – ciliated cells, basal cells, and secretory cells – as well as molecularly distinct subsets of each.

We identified several molecularly distinct, IL-13–regulated secretory cell subsets. IL-13-driven changes included the disappearance of CEACAM6^+^ only cells, and the emergence of TSPAN8^+^ and MUC5AC^+^ subsets, supporting evidence of IL-13–induced alterations in secretory cell phenotype. We confirm that transcriptional heterogeneity in secretory cells is accompanied by related changes at the protein level. Whether these subsets are functionally disparate or represent novel subsets with overlapping function, and what epigenetic processes underlie the shift in secretory cell heterogeneity require further investigation. Understanding secretory cell diversity in the context of IL-13 stimulation will be crucial in understanding asthma pathology.

Our data underline the increasing recognition of epithelial cell heterogeneity, suggesting that airway epithelial subsets may be more precisely described by sets of molecular markers rather than using traditional approaches for defining ciliated, basal and secretory cells based on morphology and/or use of more limited sets of markers (26). Traditionally, nomenclature is based principally on histologic criteria (5, 6) that fail to capture the heterogeneity evident from scRNA-seq (7–11) or antibody panels. Our protocol provides a working method for classifying airway epithelial subsets, and we expect that additional reagents can be added to this panel to further subdivide major subsets and identify other smaller populations, such as ionocytes. Furthermore, use of standard sets of molecular markers such as those developed here will promote clearer communication and allow for more meaningful comparisons across studies.

We coupled flow cytometric cell sorting with gene expression analysis and identified mRNA transcripts not detected in our scRNA-seq experiment plus several miRNAs. Although scRNA-seq is revealing airway epithelial cell transcriptomes in unprecedented resolution, current technologies have limited sensitivity and do not reliably detect low-abundance transcripts (e.g. transcription factors) that may have significant impact on cell specification and in disease. Furthermore, available scRNA-seq techniques are generally not suited for analyzing mRNA variants or small RNAs, such as miRNAs. Therefore, our sorting panel may contribute to deeper cataloging of airway epithelial subset transcriptomes. Additionally, our panels could be coupled with downstream epigenetic and proteomic analyses to further understand the specification and/or function of airway epithelial subsets in human health and disease.

There is also potential to build on the protocols we have developed. For example, although flow cytometric analysis of clinical samples was not performed, published scRNA-seq datasets suggest that the panels we developed could be useful for analyzing disaggregated cells from human airways (7, 9–11). Furthermore, although our study used a 10-parameter, 8-fluorochrome approach, application of an increased panel is viable. Future refinement and the addition of other markers for rarer epithelial subtypes (e.g. cystic fibrosis transmembrane conductance regulator (CFTR)-expressing ionocytes and doublecortin like kinase 1 (DCLK1)-expressing tuft cells (7–10)) may permit characterization of the null or minor (<1%) populations reported here. Additionally, combining the epithelial panel with existing panels for immune cells and other non-epithelial cells will permit a more comprehensive examination of lung development, airway inflammatory and immune responses, and a variety of disease states. Mass cytometry (27) or use of oligonucleotide-barcoded antibodies together with single cell sequencing (28) also promise to allow for more extensive panels of markers that could further expand our approach to increase our understanding of the function of specific subsets and their heterogeneity.

In summary, we have leveraged scRNA-seq datasets to develop flow cytometry panels for characterizing subpopulations of airway epithelial cells. These panels identified major airway epithelial cell subsets, revealed molecular heterogeneity within these populations, and permitted analysis of low abundance transcripts and miRNAs. We envisage that these panels and their future refinements will be powerful tools for interrogating airway epithelial biology.

## Supporting information

FigureE1

FigureE2

## Acknowledgements

We thank Joshua Pollack for his help analyzing Drop-seq data. We also thank the UCSF Nikon Imaging Center and Laboratory for Cell Analysis for advice and assistance with microscopy and flow cytometry, respectively.

## References

1. Maecker HT, McCoy JP, Nussenblatt R. Standardizing immunophenotyping for the Human Immunology Project. Nature Reviews Immunology 2012;12:191–200.

2. Tighe RM, Redente EF, Yu Y-R, Herold S, Sperling AI, Curtis JL, Duggan R, Swaminathan S, Nakano H, Zacharias WJ, Janssen WJ, Freeman CM, Brinkman RR, Singer BD, Jakubzick CV, Misharin AV. Improving the Quality and Reproducibility of Flow Cytometry in the Lung. An Official American Thoracic Society Workshop Report. Am J Respir Cell Mol Biol 2019;61:150–161.

3. Bonser LR, Erle DJ. The airway epithelium in asthma. Adv Immunol 2019;142:1–34.

4. Bonser LR, Erle DJ. Airway Mucus and Asthma: The Role of MUC5AC and MUC5B. J Clin Med 2017;6:.

5. Rhodin J. LXVII Ultrastructure of the Tracheal Ciliated Mucosa in Rat and Man. Annals of Otology, Rhinology & Laryngology 1959;68:964–974.

6. Widdicombe JH. Early Studies on the Surface Epithelium of Mammalian Airways. American Journal of Physiology-Lung Cellular and Molecular Physiology 2019;doi:10.1152/ajplung.00240.2019.

7. Plasschaert LW, Žilionis R, Choo-Wing R, Savova V, Knehr J, Roma G, Klein AM, Jaffe AB. A single-cell atlas of the airway epithelium reveals the CFTR-rich pulmonary ionocyte. Nature 2018;560:377.

8. Montoro DT, Haber AL, Biton M, Vinarsky V, Lin B, Birket SE, Yuan F, Chen S, Leung HM, Villoria J, Rogel N, Burgin G, Tsankov AM, Waghray A, Slyper M, Waldman J, Nguyen L, Dionne D, Rozenblatt-Rosen O, Tata PR, Mou H, Shivaraju M, Bihler H, Mense M, Tearney GJ, Rowe SM, Engelhardt JF, Regev A, Rajagopal J. A revised airway epithelial hierarchy includes CFTR-expressing ionocytes. Nature 2018;560:319.

9. Ordovas-Montanes J, Dwyer DF, Nyquist SK, Buchheit KM, Vukovic M, Deb C, Wadsworth MH, Hughes TK, Kazer SW, Yoshimoto E, Cahill KN, Bhattacharyya N, Katz HR, Berger B, Laidlaw TM, Boyce JA, Barrett NA, Shalek AK. Allergic inflammatory memory in human respiratory epithelial progenitor cells. Nature 2018;560:649.

10. Duclos GE, Teixeira VH, Autissier P, Gesthalter YB, Reinders-Luinge MA, Terrano R, Dumas YM, Liu G, Mazzilli SA, Brandsma C-A, Berge M van den, Janes SM, Timens W, Lenburg ME, Spira A, Campbell JD, Beane J. Characterizing smoking-induced transcriptional heterogeneity in the human bronchial epithelium at single-cell resolution. Science Advances 2019;5:eaaw3413.

11. García SR, Deprez M, Lebrigand K, Cavard A, Paquet A, Arguel M-J, Magnone V, Truchi M, Caballero I, Leroy S, Marquette C-H, Marcet B, Barbry P, Zaragosi L-E. Novel dynamics of human mucociliary differentiation revealed by single-cell RNA sequencing of nasal epithelial cultures. Development 2019;146:.

12. Maestre-Batlle D, Pena OM, Hirota JA, Gunawan E, Rider CF, Sutherland D, Alexis NE, Carlsten C. Novel flow cytometry approach to identify bronchial epithelial cells from healthy human airways. Sci Rep 2017;7:1–9.

13. Griggs TF, Bochkov YA, Basnet S, Pasic TR, Brockman-Schneider RA, Palmenberg AC, Gern JE. Rhinovirus C targets ciliated airway epithelial cells. Respir Res 2017;18:84.

14. Rock JR, Onaitis MW, Rawlins EL, Lu Y, Clark CP, Xue Y, Randell SH, Hogan BLM. Basal cells as stem cells of the mouse trachea and human airway epithelium. PNAS 2009;106:12771–12775.

15. Koh KD, Siddiqui S, Cheng D, Bonser LR, Sun DI, Zlock LT, Finkbeiner WE, Woodruff PG, Erle DJ. Efficient RNP-Directed Human Gene Targeting Reveals SPDEF is Required for IL-13-Induced Mucostasis. Am J Respir Cell Mol Biol 2019;doi:10.1165/rcmb.2019-0266OC.

16. Fulcher ML, Gabriel S, Burns KA, Yankaskas JR, Randell SH. Well-Differentiated Human Airway Epithelial Cell Cultures. In: Picot J, editor. Human Cell Culture Protocols Totowa, NJ: Humana Press; 2005. p. 183–206.doi:10.1385/1-59259-861-7:183.

17. Whitcutt MJ, Adler KB, Wu R. A biphasic chamber system for maintaining polarity of differentiation of cultured respiratory tract epithelial cells. In Vitro Cell Dev Biol 1988;24:420–428.

18. Bonser LR, Zlock L, Finkbeiner W, Erle DJ. Epithelial tethering of MUC5AC-rich mucus impairs mucociliary transport in asthma. J Clin Invest 2016;126:2367–2371.

19. Macosko EZ, Basu A, Satija R, Nemesh J, Shekhar K, Goldman M, Tirosh I, Bialas AR, Kamitaki N, Martersteck EM, Trombetta JJ, Weitz DA, Sanes JR, Shalek AK, Regev A, McCarroll SA. Highly Parallel Genome-wide Expression Profiling of Individual Cells Using Nanoliter Droplets. Cell 2015;161:1202–1214.

20. Lukinavičius G, Reymond L, D’Este E, Masharina A, Göttfert F, Ta H, Güther A, Fournier M, Rizzo S, Waldmann H, Blaukopf C, Sommer C, Gerlich DW, Arndt H-D, Hell SW, Johnsson K. Fluorogenic probes for live-cell imaging of the cytoskeleton. Nature Methods 2014;11:731–733.

21. Oltean A, Schaffer AJ, Bayly PV, Brody SL. Quantifying Ciliary Dynamics during Assembly Reveals Stepwise Waveform Maturation in Airway Cells. Am J Respir Cell Mol Biol 2018;59:511–522.

22. Christenson SA, van den Berge M, Faiz A, Inkamp K, Bhakta N, Bonser LR, Zlock LT, Barjaktarevic IZ, Barr RG, Bleecker ER, Boucher RC, Bowler RP, Comellas AP, Curtis JL, Han MK, Hansel NN, Hiemstra PS, Kaner RJ, Krishnanm JA, Martinez FJ, O’Neal WK, Paine R, Timens W, Wells JM, Spira A, Erle DJ, Woodruff PG. An airway epithelial IL-17A response signature identifies a steroid-unresponsive COPD patient subgroup. J Clin Invest 2019;129:169–181.

23. Pan J, Adair-Kirk TL, Patel AC, Huang T, Yozamp NS, Xu J, Reddy EP, Byers DE, Pierce RA, Holtzman MJ, Brody SL. Myb permits multilineage airway epithelial cell differentiation. Stem Cells 2014;32:3245–3256.

24. Song R, Walentek P, Sponer N, Klimke A, Lee JS, Dixon G, Harland R, Wan Y, Lishko P, Lize M, Kessel M, He L. miR-34/449 miRNAs are required for motile ciliogenesis by repressing cp110. Nature 2014;510:115–120.

25. Biton M, Levin A, Slyper M, Alkalay I, Horwitz E, Mor H, Kredo-Russo S, Avnit-Sagi T, Cojocaru G, Zreik F, Bentwich Z, Poy MN, Artis D, Walker MD, Hornstein E, Pikarsky E, Ben-Neriah Y. Epithelial microRNAs regulate gut mucosal immunity via epithelium–T cell crosstalk. Nature Immunology 2011;12:239–246.

26. Bonser LR, Erle DJ. Putting Mucins on the Map. American Journal of Respiratory and Critical Care Medicine 2018;doi:10.1164/rccm.201809-1818ED.

27. Bandura DR, Baranov VI, Ornatsky OI, Antonov A, Kinach R, Lou X, Pavlov S, Vorobiev S, Dick JE, Tanner SD. Mass cytometry: technique for real time single cell multitarget immunoassay based on inductively coupled plasma time-of-flight mass spectrometry. Anal Chem 2009;81:6813–6822.

28. Akkaya B, Miozzo P, Holstein AH, Shevach EM, Pierce SK, Akkaya M. A Simple, Versatile Antibody-Based Barcoding Method for Flow Cytometry. The Journal of Immunology 2016;197:2027–2038.

29. Piperno G, Fuller MT. Monoclonal antibodies specific for an acetylated form of alpha-tubulin recognize the antigen in cilia and flagella from a variety of organisms. J Cell Biol 1985;101:2085–2094.

30. Basnet S, Bochkov YA, Brockman-Schneider RA, Kuipers I, Aesif SW, Jackson DJ, Lemanske RF, Ober C, Palmenberg AC, Gern JE. CDHR3 Asthma-Risk Genotype Affects Susceptibility of Airway Epithelium to Rhinovirus C Infections. Am J Respir Cell Mol Biol 2019;61:450–458.

31. Bønnelykke K, Sleiman P, Nielsen K, Kreiner-Møller E, Mercader JM, Belgrave D, den Dekker HT, Husby A, Sevelsted A, Faura-Tellez G, Mortensen LJ, Paternoster L, Flaaten R, Mølgaard A, Smart DE, Thomsen PF, Rasmussen MA, Bonàs-Guarch S, Holst C, Nohr EA, Yadav R, March ME, Blicher T, Lackie PM, Jaddoe VWV, Simpson A, Holloway JW, Duijts L, Custovic A, et al. A genome-wide association study identifies CDHR3 as a susceptibility locus for early childhood asthma with severe exacerbations. Nat Genet 2014;46:51–55.

32. Beane J, Sebastiani P, Liu G, Brody JS, Lenburg ME, Spira A. Reversible and permanent effects of tobacco smoke exposure on airway epithelial gene expression. Genome Biol 2007;8:R201.

33. Ooi AT, Gower AC, Zhang KX, Vick JL, Hong L, Nagao B, Wallace WD, Elashoff DA, Walser TC, Dubinett SM, Pellegrini M, Lenburg ME, Spira A, Gomperts BN. Molecular Profiling of Premalignant Lesions in Lung Squamous Cell Carcinomas Identifies Mechanisms Involved in Stepwise Carcinogenesis. Cancer Prev Res 2014;7:487–495.

34. Shikotra A, Choy DF, Siddiqui S, Arthur G, Nagarkar DR, Jia G, Wright AKA, Ohri CM, Doran E, Butler CA, Hargadon B, Abbas AR, Jackman J, Wu LC, Heaney LG, Arron JR, Bradding P. A CEACAM6-High Airway Neutrophil Phenotype and CEACAM6-High Epithelial Cells Are Features of Severe Asthma. J Immunol 2017;198:3307–3317.

