## Supplementary figures and images for "Flow cytometric analysis and purification of airway epithelial cell subsets"

### FigureE1

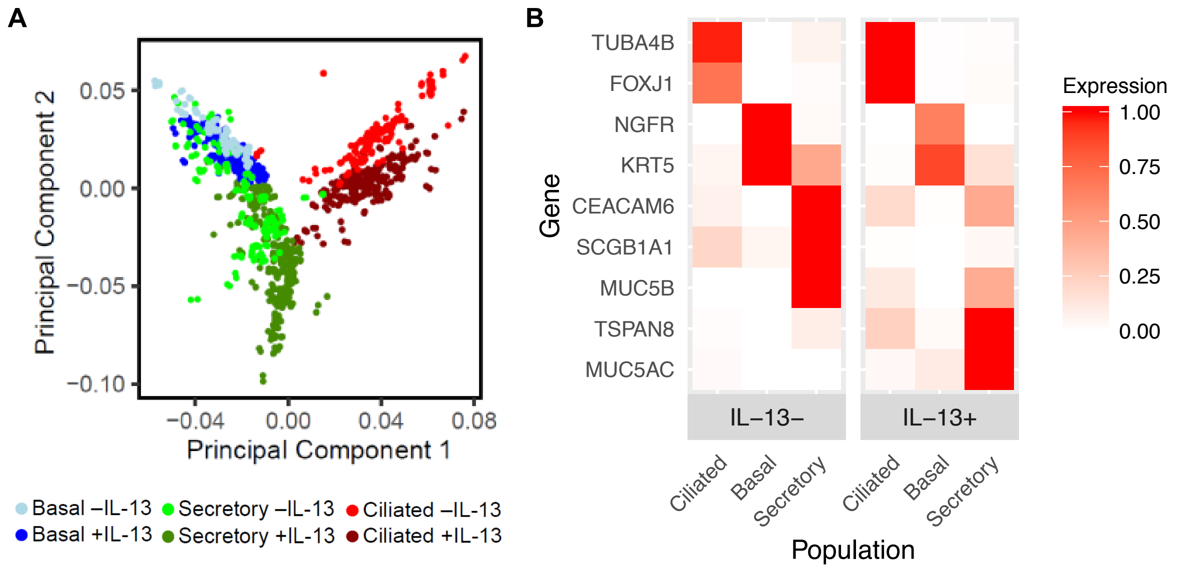

### FigureE2

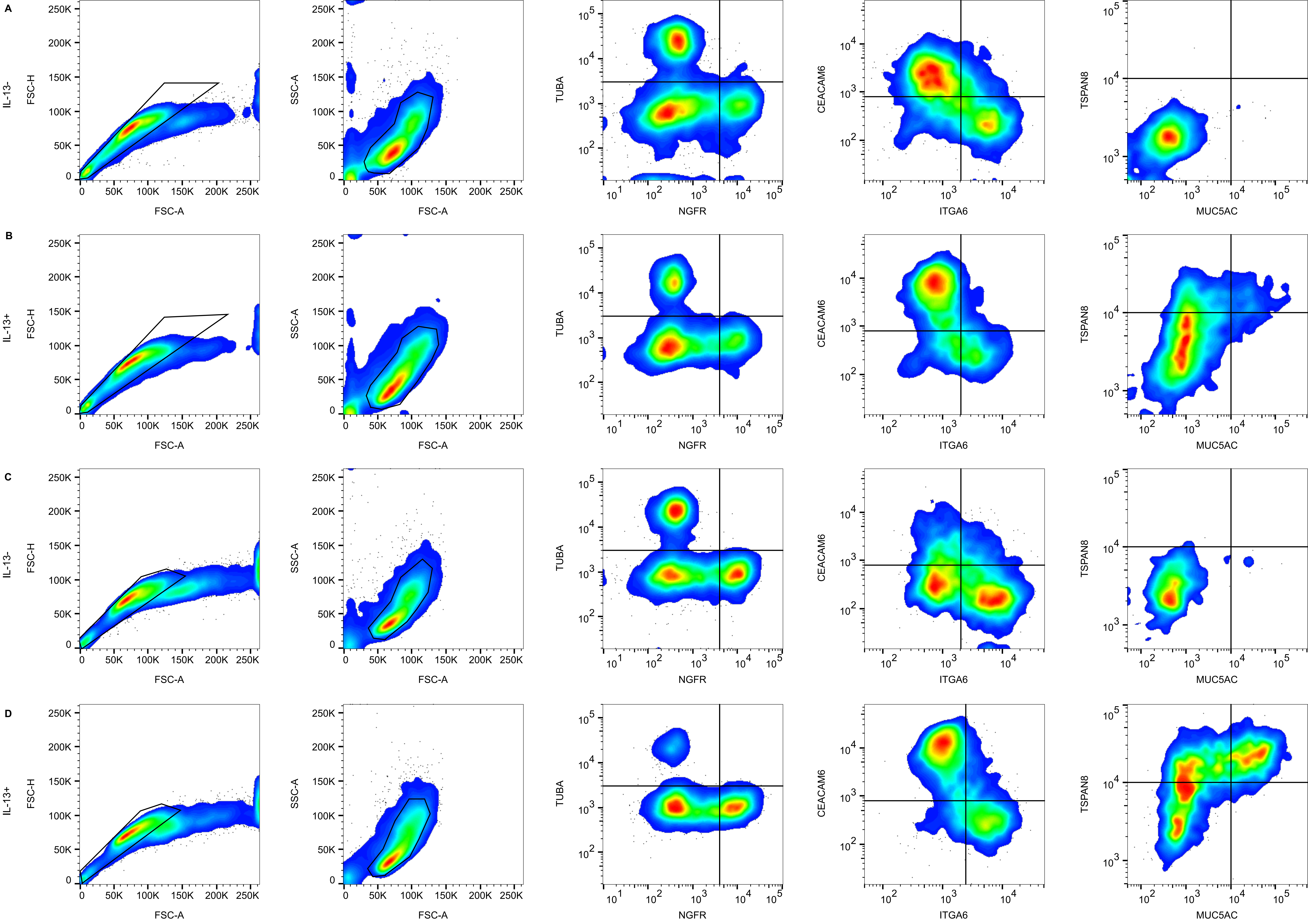
