## Supplementary material for "Flow cytometric analysis and purification of airway epithelial cell subsets": Online data supplement

^1^Lung Biology Center, ^2^Cardiovascular Research Institute, ^3^Department of Microbiology and Immunology, ^4^Division of Pulmonary, Critical Care, Sleep, and Allergy, ^5^Sandler Asthma Basic Research Center, ^6^Department of Bioengineering and Therapeutic Sciences, ^7^Department of Pathology, and ^8^UCSF CoLabs, University of California San Francisco, San Francisco, CA 94143 and ^9^Department of Respiratory and Critical Care Medicine, Renmin Hospital of Wuhan University, Wuhan, China 430060

**Supplementary Methods**

**Human bronchial epithelial cell (HBEC) culture**

HBECs were isolated from lungs not used for transplantation (n=12; written consent was not required, as materials were leftover clinical samples obtained from deidentified individuals) and cultured on 12-mm Transwell inserts (Corning, Corning, NY) at air-liquid interface (ALI) as described previously (1, 2). After 16 days in culture at ALI, cultures were supplemented with IL-13 (10 ng/mL, Peprotech, Rocky Hill, NJ) for 7 days to induce goblet cell production (3).

**Generating single cell suspensions from HBECs**

Accumulated mucus was removed by washing the apical surface of cultures in 10 mM dithiothrietol (DTT; ThermoFisher Scientific, Fremont, CA, USA) in phosphate buffered saline (PBS; Life Technologies, Carlsbad, CA) for 10 minutes. The apical surface was then washed three times in PBS to remove excess DTT. Basolateral media was aspirated, the basolateral membrane rinsed briefly in 1 mL PBS, and cell suspensions generated by incubating cells both apically and basolaterally in 0.25 % trypsin-EDTA (ThermoFisher Scientific) for 10-15 minutes at 37 °C. The trypsinized cells were neutralized in DMEM/F-12 containing 5% FBS, passed through a 100-µm filter, and counted using a hemocytometer.

**Drop-seq**

Drop-seq libraries were prepared from HBEC suspensions as previously described (4). Briefly, using microfluidics, we created droplets containing a single cell and a distinctly barcoded bead. Within each droplet, the cells were lysed releasing their mRNAs that hybridized to primers on the bead surface. We then broke the droplets, collected the beads, and reverse transcribed the mRNAs into cDNAs, generating a set of beads called STAMPS (single cell transcriptomes attached to microparticles). We then amplified the barcoded STAMPS in bulk and performed paired-end RNA sequencing. We aligned each mRNA to the genome to identify the gene of origin and used the STAMP barcode to identify the cell of origin. We clustered cells using known markers for airway epithelial subpopulations. For the purpose of identifying novel markers of major subsets, we selected cells expressing combinations of known markers for basal cells *(*e.g., *NGFR, ITGA6, and KRT5)*, secretory cells (*SCGB1A1*, *MUC5B*, *MUC5AC*), and ciliated cells (*FOXJ1)* and identified other genes that were differentially expressed between these subsets.

**Antibody conjugation**

For flow cytometry analysis and cell sorting, CDHR3 and MUC5AC antibodies were conjugated in house. Mouse monoclonal anti-MUC5AC was concentrated to 1 mg/mL using the Antibody Clean Up and Concentrator kit. Rabbit polyclonal anti-CDHR3 was isolated using AbPure immunomagnetic beads to remove glycerol from the antibody buffer, eluted, and concentrated as per MUC5AC. Concentrated, purified MUC5AC and CDHR3 antibodies were conjugated to APC-Cy7 and R-PE, respectively, using Lightning Link Rapid conjugation kits (Abcam, Cambridge, UK) as per the manufacturer’s instructions.

**Flow cytometric analysis**

For flow cytometric analysis, single cell HBEC suspensions were fixed in 0.5% paraformaldehyde (PFA; ThermoFisher Scientific). PFA-fixed cells were washed twice in PBS containing protease inhibitors (PI) and sodium butyrate; after the second wash, excess buffer was removed leaving ~30 uL volume, and cells were frozen at -80 °C if not used immediately. Prior to staining, cells were thawed if necessary and resuspended in blocking buffer (5% [v/v] normal goat serum [Jackson ImmunoResearch Inc., West Grove, PA] in PBS) on a rotator for 15 min to block non-specific binding. Appropriate volumes of cell surface marker antibodies (see Table S1) were placed in 96-well plates, and blocked cells were added; cells were incubated at 4 °C for 30 minutes on a rocker. Plates were centrifuged and washed twice in blocking buffer. Cells were subsequently permeabilized in blocking buffer containing 0.2% (w/v) saponin (MilliporeSigma) for 15 minutes. Permeabilized cells were then transferred to plates containing intracellular marker antibodies and incubated at 4 °C for 30 minutes on a rocker. Cells were then washed twice and resuspended in eBioScience Flow Cytometry Staining Buffer (ThermoFisher Scientific) and transferred to pre-labelled tubes for flow cytometric analysis.

Data were acquired on a BD FACS CantoII flow cytometer using BD FACS Diva software (BD Biosciences), and compensation and data analyses were performed using FlowJo software (TreeStar, Ashland, OR). Prior to analysis of subset staining, doublets were omitted from the analysis by plotting forward scatter height (FSC-H) against forward scatter area (FSC-A) and gating out events for which the FSC-A was larger than expected for the FSC-H. Subsequently, debris, identified as having very low side scatter area (SSC-A) and FSC-A, was also removed from the analysis. Isotype controls were used to assess positive antibody staining: gates were placed at the 99^th^ percentile of intensity seen with the isotype control to identify cells that had specific staining in the channel of interest. The gating strategy described in the text was then used to identify the relevant HBEC subpopulations. The “analytical panel” is provided in Table E1, and a detailed step-by-step method provided in the Supplementary Document.

**Flow cytometric cell sorting**

To identify ciliated cells from live cultures, we utilized SiR-tubulin (Spirochrome/Cytoskeleton Inc., Denver, CO): a live cell dye which stains microtubules, a major structural component of cilia (6). SiR-tubulin was prepared as per the manufacturer’s instructions and diluted 1:1000 in ALI media at a final concentration of 1 µM; medium was also supplemented with 10 µM verapamil, a broad spectrum efflux pump inhibitor, reported to greatly improve SiR-tubulin staining (6). HBEC cultures were incubated with SiR-tubulin staining media for 1 hour at 37 °C in a humidified atmosphere containing 5% CO_2_. Cultures were then washed with PBS and trypsinized to generate single cell HBEC suspensions. HBECs were centrifuged and resuspended in fixable viability dye eFluor 450 (ThermoFisher Scientific). Cells were then blocked in blocking buffer supplemented with RNase I on a rotator for 15 minutes. Cell sorting was performed on a BD FACS Aria II instrument (BD Biosciences) with a similar configuration as the CantoII; compensation was performed with UltraComp eBeads (Invitrogen) automatically within BD FACS Diva prior to acquisition. Singlets were discriminated from doublets using FSC-H vs FSC-A and debris (low SSC-A and FSC-A) was also excluded. Isotype controls were used to assess nonspecific staining: gates were placed at the 99^th^ percentile of the isotype control to identify cells that had specific staining in the channel of interest. Cell populations were then identified using the sequential gating strategy described in the text. Cells were sorted and collected in cation-free PBS containing HEPES and BSA. The sorting panel is shown in Table E1, and a detailed step-by-step method is provided in the Supplementary Document.

**Preparation of cells for cytospin**

HBECs were fixed in 0.5% paraformaldehyde (PFA; Electron Microscopy Sciences, Fort Washington, PA) for 8-10 minutes at room temperature, and deposited on microscope slides by cytospin as previously described (7). The fixed dried cells were permeabilized in 0.1% (v/v) Triton X100 for 20 min and blocked in either 5% (v/v) normal goat serum or donkey serum (according to the origin of the primary antibody; Jackson ImmunoResearch Inc., West Grove, PA) in phosphate buffered saline with 0.05% Tween 20 for 30 minutes at room temperature prior to primary antibody incubation. The primary antibodies used are listed in Table S1.

After overnight primary antibody incubation at 4 °C, slides were washed and incubated with appropriate secondary antibodies (see Table S1). All secondary antibodies (Jackson ImmunoResearch Inc.) were used at a 1:200 dilution and incubations were performed at room temperature for 2-3 h. Simultaneously, 4',6-diamidino-2-phenylindole (DAPI) was used to counterstain nuclei. Slides were washed with PBS and post-fixed in 2% (v/v) PFA and mounted with Fluoromount-G (Southern Biotech, Birmingham, AL). Images were acquired using a Yokagawa CSU22 spinning-disk confocal microscope connected to a Nikon Ti-E (Nikon Imaging Center, UCSF). Slides were placed on the microscope stage and images were acquired using appropriate lasers and a 60× oil objective. Identical acquisition settings were used throughout acquisition.

**RNA Isolation**

Following live sorting, HBECs were lysed in Buffer and homogenized by vortexing. Total RNA was isolated using the Rneasy Kit (Qiagen; Hilden, Germany) according to the manufacturer’s instructions and used for mRNA and miRNA analysis.

**mRNA quantification**

mRNA was reverse-transcribed using the SuperScript III First-Strand Synthesis System (ThermoFisher Scientific), and the resulting cDNA was analyzed by quantitative real-time PCR (qRT-PCR) using PowerUp SYBR Green (ThermoFisher Scientific); primer sequences are presented in Table E2. The mean value of three technical replicates was used for analysis. From our Drop-seq dataset, we identified several genes that showed little variance across the sorted subsets and assessed their expression by qRT-PCR. *SCAF11* showed the least variance across samples, and mRNA levels were subsequently normalized to *SCAF11* expression. Comparisons were made using the using ΔΔC_t_ method (8).

**miRNA quantification**

Reverse transcription was performed using miRNA-specific stem-loop primers and MultiScribe Reverse Transcriptase (ThermoFisher Scientific). cDNA was pre-amplified using AmpliTaq Gold DNA Polymerase (ThermoFisher Scientific) and then purified using ExoSAP-IT (ThermoFisher Scientific) followed by G-50 columns (GE Healthcare, Chicago, IL). The qRT-PCR assay was performed on a ViiA 7 Real-Time PCR system (ThermoFisher Scientific) using TaqMan Universal PCR Master Mix (ThermoFisher Scientific). All primers and TaqMan probes are listed in Table S2. miRNA expression was normalized to hsa-miR-103-3p and hsa-miR-191-5p expression (9) and comparisons made using the using ΔΔC_t_ method (8).

**Statistical analysis**

Flow cytometric analysis was performed using FlowJo. IL-13 effects on individual marker staining was analyzed using the Student’s t-test in R, and figures were produced using ggplot2 (10). For TSNE analysis, samples from each donor grown in the absence or presence of IL-13 were downsampled to 3000 cells so that each made an equal contribution to the final analysis and concatenated within FlowJo. For analysis of cell heterogeneity, each cell was characterized by the presence or absence of staining for each of the eight markers in the analysis panel (2^8^ = 256 potential subsets). Subsets containing >1% of total cells and present in all three donors in either unstimulated or IL-13-stimulated cultures were used included in subsequent analyses. The effect of IL-13 on HBEC subsets was analyzed by Student’s t-test in R (10). qRT-PCR data was analyzed in R. Figures were produced using ggplot2 (10).

**Supplementary References**

1.	Fulcher ML, Gabriel S, Burns KA, Yankaskas JR, Randell SH. Well-Differentiated Human Airway Epithelial Cell Cultures. In: Picot J, editor. *Human Cell Culture Protocols* Totowa, NJ: Humana Press; 2005. p. 183–206.doi:10.1385/1-59259-861-7:183.

2.	Whitcutt MJ, Adler KB, Wu R. A biphasic chamber system for maintaining polarity of differentiation of cultured respiratory tract epithelial cells. *In Vitro Cell Dev Biol* 1988;24:420–428.

3.	Bonser LR, Zlock L, Finkbeiner W, Erle DJ. Epithelial tethering of MUC5AC-rich mucus impairs mucociliary transport in asthma. *J Clin Invest* 2016;126:2367–2371.

4.	Macosko EZ, Basu A, Satija R, Nemesh J, Shekhar K, Goldman M, Tirosh I, Bialas AR, Kamitaki N, Martersteck EM, Trombetta JJ, Weitz DA, Sanes JR, Shalek AK, Regev A, McCarroll SA. Highly Parallel Genome-wide Expression Profiling of Individual Cells Using Nanoliter Droplets. *Cell* 2015;161:1202–1214.

5.	Schindelin J, Arganda-Carreras I, Frise E, Kaynig V, Longair M, Pietzsch T, Preibisch S, Rueden C, Saalfeld S, Schmid B, Tinevez J-Y, White DJ, Hartenstein V, Eliceiri K, Tomancak P, Cardona A. Fiji: an open-source platform for biological-image analysis. *Nat Methods* 2012;9:676–682.

6.	Lukinavičius G, Reymond L, D’Este E, Masharina A, Göttfert F, Ta H, Güther A, Fournier M, Rizzo S, Waldmann H, Blaukopf C, Sommer C, Gerlich DW, Arndt H-D, Hell SW, Johnsson K. Fluorogenic probes for live-cell imaging of the cytoskeleton. *Nature Methods* 2014;11:731–733.

7.	Koh CM. Chapter Sixteen - Preparation of Cells for Microscopy using Cytospin. In: Lorsch J, editor. *Methods in Enzymology* Academic Press; 2013. p. 235–240.

8.	Livak KJ, Schmittgen TD. Analysis of Relative Gene Expression Data Using Real-Time Quantitative PCR and the 2−ΔΔCT Method. *Methods* 2001;25:402–408.

9.	Peltier HJ, Latham GJ. Normalization of microRNA expression levels in quantitative RT-PCR assays: identification of suitable reference RNA targets in normal and cancerous human solid tissues. *RNA* 2008;14:844–852.

10.	Wickham H. *ggplot2: Elegant Graphics for Data Analysis*. Springer; 2016.

**Table E1: Antibody details**

| **Analytical panel** |  |  |  |  |  |  |  |  |
| --- | --- | --- | --- | --- | --- | --- | --- | --- |
| Antigen | Gene | Fluorophore | Clone | Isotype | Source | Ab/test (µL) | Location* | Target cell |
| TUBA | TUBA1A1 etc. | AF647 | 6-11B-1 | mIgG2b | Santa Cruz | 1 | IC | Ciliated |
| CDHR3 | CDHR3 | R-PE | HPA011218 | rpAb | Sigma | 1 | CSM | Ciliated |
| CD49f | ITGA6 | PE-CF594 | GoH3 | ratIgG2a | BD | 0.25 | CSM | Basal |
| CD271 | NGFR | PE Cy7 | ME20.4 | mIgG1 | Biolegend | 0.1 | CSM | Basal |
| CD66c | CEACAM6 | BV510 | B6.2 | mIgG1 | BD | 1 | CSM | Secretory |
| CD66e | CEACAM5 | APC | 487609 | mIgG2a | R&D | 1 | CSM | Secretory |
| TSPAN8 | TSPAN8 | AF405 | FAB4734V | ratIgG2b | R&D | 1.5 | CSM | Goblet |
| MUC5AC | MUC5AC | APC Cy7 | 45M1 | mIgG1 | Thermo | 0.25 | IC | Goblet |
| **Sorting panel** |  |  |  |  |  |  |  |  |
| Stain/Antigen | Gene | Fluorophore | Clone | Isotype | Source | Ab/test (uL) | Target cell |  |
| SiR-tubulin | TUBA | APC | n/a | n/a | Cytoskeleton, Inc. | 0.05 | Ciliated |  |
| Fixable Viability Dye | - | eFluor450 | n/a | n/a | Invitrogen | 0.5 | Live cells |  |
| CD271 | NGFR | PE Cy7 | ME20.4 | mIgG1 | Biolegend | 0.1 | Basal |  |
| CD66c | CEACAM6 | BV510 | B6.2 | mIgG1 | BD | 1 | Secretory |  |
| TSPAN8 | TSPAN8 | FITC | FAB4734V | ratIgG2b | R&D | 0.5 | Goblet |  |

* IC, intracellular; CSM, cell surface (membrane)

**Table E2: Primer sequences**

| Name | Sequence |
| --- | --- |
| q.TUBA1A.2 | AACTATGCCCGAGGGCACT |
| q.TUBA1A.3 | AGAAGCCCTGGAGACCCG |
| q.CDHR3.1 | TGAGGCCTTCGATCCAGAAG |
| q.CDHR3.2 | GGTGCCATTAGCAGACATTCTG |
| q.FOXJ1.1 | CGAGGCACTTTGATGAAGC |
| q.FOXJ1.3 | ACAAGTGGATCACGGACAACTT |
| q.MYB.1 | GCCAGCCCACTGTTAACAAC |
| q.MYB.2 | ACAGGGTATGGAACATGACTGG |
| q.NGFR.3 | AGCTCCGCGAGTGCACAC |
| q.NGFR.4 | GGTGTGGACCGTGTAATCCAA |
| q.ITGA6.1 | CCCGCTGGTTATAATCCTTCA |
| q.ITGA6.2 | GAACTCTTGAGGATAGCCCAGAT |
| q.KRT5.3 | AGCAGTGGTACGCTTGTTGATT |
| q.KRT5.4 | GCCTGGACTCAGAGCTGAGAA |
| q.TSLP.1 | GCTATCTGGTGCCCAGGCTAT |
| q.TSLP.2 | CGACGCCACAATCCTTGTAA |
| q.CEACAM6.1 | CTGGCCTCAATAGGACCACAGT |
| q.CEACAM6.2 | GCCAGCACTCCAATCGTGAT |
| q.SCGB1A1.3 | GCTGAAGAAGCTGGTGGACAC |
| q.SCGB1A1.4 | TGCTAATTACACAGTGAGCTTTGG |
| q.MUC5B.3 | GCTCCAAGGCCATCAAGCT |
| q.MUC5B.4 | GGTCTCGATGACCAGGAAGATC |
| q.CEACAM5.1 | GGTGCATCCCCTGGCAGA |
| q.CEACAM5.2 | GCTTGGCAGTGGTGGGC |
| q.TSPAN8.1 | CAATCAGCAGCTCCATTGAC |
| q.TSPAN8.2 | TTCCAGGAAGCCATAATTGTG |
| q.MUC5AC.1 | CAACATCAGGAACAGCTTCGA |
| q.MUC5AC.2 | GAGCACCAGTGCTGAGCAT |
| q.SPDEF.1 | AAGTTGGCACTGCAGCAGAC |
| q.SPDEF.2 | GGGGATACGCTGCTCAGAC |
| q.PRB1.1 | GCCTCCCCAGTCATCTAGGA |
| q.PRB1.2 | CCAATGTCATGGAATTTGAATCA |
| q.SCAF11.1 | CCCTTTTCCAAAGAGAAATGAAG |
| q.SCAF11.2 | CTCACTGTACAACAGACCAGTGG |
| hsa-miR-34c-5p F | ACACTCCAGCTGGGAGGCAGTGTAGTTAGCT |
| hsa-miR-34c-5p R | CTCAACTGGTGTCGTGGAGTCGGCAATTCAGTTGAGGCAATCAG |
| hsa-miR-34c-5p P | TTCAGTTGAGGCAATCAG |
| hsa-miR-449a F | ACACTCCAGCTGGGTGGCAGTGTATTGTTA |
| hsa-miR-449a R | CTCAACTGGTGTCGTGGAGTCGGCAATTCAGTTGAGACCAGCTA |
| hsa-miR-449a P | TTCAGTTGAGACCAGCTA |
| hsa-miR-375 F | ACACTCCAGCTGGGTTTGTTCGTTCGGCTC |
| hsa-miR-375 R | CTCAACTGGTGTCGTGGAGTCGGCAATTCAGTTGAGTCACGCGA |
| hsa-miR-375 P | TTCAGTTGAGTCACGCGA |
| hsa-miR-103a-3p F | ACACTCCAGCTGGGAGCAGCATTGTACAGGG |
| hsa-miR-103a-3p R | CTCAACTGGTGTCGTGGAGTCGGCAATTCAGTTGAGTCATAGCC |
| hsa-miR-103a-3p P | TTCAGTTGAGTCATAGCC |

**Table E3: Molecular heterogeneity of FACS identified subsets***

| TUBA | CDHR3 | NGFR | ITGA6 | CEACAM6 | CEACAM5 | TSPAN8 | MUC5AC | A_UN | B_UN | C_UN | Mean_UN | A_13 | B_13 | C_13 | Mean_13 |
| --- | --- | --- | --- | --- | --- | --- | --- | --- | --- | --- | --- | --- | --- | --- | --- |
| – | – | + | + | – | – | – | – | 19.3% | 13.3% | 19.1% | 17.2% | 21.7% | 14.8% | 18.8% | 18.4% |
| + | + | – | – | – | – | – | – | 16.2% | 12.0% | 15.1% | 14.4% | 2.5% | 1.6% | 2.3% | 2.1% |
| – | – | – | – | + | + | – | – | 4.3% | 21.6% | 9.7% | 11.9% | 3.5% | 14.0% | 6.0% | 7.9% |
| – | – | – | + | – | – | – | – | 13.5% | 8.8% | 7.7% | 10.0% | 7.5% | 4.9% | 2.3% | 4.9% |
| – | – | – | – | – | – | – | – | 9.0% | 6.8% | 10.9% | 8.9% | 2.4% | 9.8% | 11.1% | 7.8% |
| – | – | – | – | + | – | – | – | 6.8% | 5.1% | 13.2% | 8.4% | 0.0% | 0.0% | 0.0% | 0.0% |
| + | + | – | – | + | – | – | – | 2.2% | 7.7% | 6.0% | 5.3% | 3.9% | 3.3% | 2.0% | 3.1% |
| + | – | – | – | – | – | – | – | 3.7% | 1.7% | 2.7% | 2.7% | 0.0% | 0.0% | 1.3% | 0.4% |
| – | + | + | + | – | – | – | – | 0.0% | 4.3% | 2.2% | 2.2% | 9.4% | 3.1% | 0.0% | 4.1% |
| – | – | – | – | – | + | – | – | 0.0% | 2.3% | 1.5% | 1.2% | 1.6% | 6.0% | 6.6% | 4.7% |
| + | + | – | – | + | – | – | + | 0.0% | 1.2% | 0.0% | 0.4% | 2.3% | 2.9% | 1.6% | 2.3% |
| + | + | – | – | + | + | – | – | 0.0% | 1.1% | 0.0% | 0.4% | 2.5% | 2.0% | 1.3% | 1.9% |
| – | – | – | – | + | + | + | – | 0.0% | 0.0% | 0.0% | 0.0% | 6.3% | 8.1% | 8.2% | 7.6% |
| – | + | – | – | + | + | + | + | 0.0% | 0.0% | 0.0% | 0.0% | 10.6% | 1.9% | 4.6% | 5.7% |
| – | – | – | – | + | + | + | + | 0.0% | 0.0% | 0.0% | 0.0% | 4.0% | 4.6% | 7.8% | 5.5% |
| + | + | – | – | + | + | – | + | 0.0% | 0.0% | 0.0% | 0.0% | 3.1% | 3.8% | 1.8% | 2.9% |
| – | – | – | – | – | + | + | – | 0.0% | 0.0% | 0.0% | 0.0% | 1.2% | 1.6% | 3.3% | 2.1% |
| – | + | – | – | + | + | + | – | 0.0% | 0.0% | 0.0% | 0.0% | 1.9% | 1.3% | 1.4% | 1.5% |
| Other (sum of all subsets that represent <1% of cells in both unstimulated and IL-13-stimulated cultures) | | | | | | | | 25.0% | 14.0% | 12.0% | 17.0% | 15.6% | 16.2% | 19.6% | 17.1% |

* A, B, and C represent cultures from three different donors; UN = unstimulated, 13 = IL-13 stimulated

**Supplemental Figure Legends**

**Figure E1. Drop-seq analysis of HBEC cultures.** Unstimulated or IL-13-stimulated HBECs were trypsinized and single cells subjected to scRNA-sequencing using Drop-seq. (*A*) Principal component analysis revealed three major cell clusters corresponding to basal, ciliated, and secretory cells; basal cells largely overlapped irrespective of treatment, however, for secretory and ciliated cells, two clusters separated by IL-13 treatment were observed. (*B*) Heatmap illustrating select cell-specific transcripts (defined as genes more highly expressed in one cell type than the others (FDR < 5%)).

**Figure E2. Gating strategy for analytical panel in two additional donors.** The same gating strategy employed in Figure 2M-Q was applied to cells from two other donors. Debris and doublets were removed and singlets were gated on NGFR and TUBA. To identify goblet cells, IL-13-stimulated TUBA-NGFR- singlets were gated on ITGA6 and CEACAM5. CEACAM5+ cells were then gated on MUC5AC and TSPAN8. (*A*) Donor A, unstimulated. (*B*) Donor A, IL-13-stimulated. (*C*) Donor B, unstimulated. (*D*) Donor B, IL-13-stimulated.
