## Supplementary material for "Flow cytometric analysis and purification of airway epithelial cell subsets": Online supplementary document

**Online supplementary document: Detailed HBEC flow cytometry protocols, version 20200215**

^1^Lung Biology Center, ^2^Cardiovascular Research Institute, ^3^Department of Microbiology and Immunology, ^4^Division of Pulmonary, Critical Care, Sleep, and Allergy, ^5^Sandler Asthma Basic Research Center, ^6^Department of Bioengineering and Therapeutic Sciences, ^7^Department of Pathology, and ^8^UCSF CoLabs, University of California San Francisco, San Francisco, CA 94143 and ^9^Department of Respiratory and Critical Care Medicine, Renmin Hospital of Wuhan University, Wuhan, China 430060

**Generating HBE single cell suspension**

1. Prewarm solutions at 37°C.
2. Remove fully differentiated HBE cultures (maintained at ALI for ~3 weeks) from the incubator and wash the apical surface with ~1.5 µL/mm^2^ of culture area (1) (~75 µL/6.5mm transwell; ~150 µL/12mm transwell) PBS containing 10 mM Dithiothrietol (DTT, MilliporeSigma, St Luois, MO) for 10 minutes at 37 °C to remove accumulated mucus (Note 1). Carefully remove the DTT wash so as not to disturb the underlying epithelium (Note 2).
3. Wash the apical surface of the cultures with ~1.5 µL PBS/mm^2^ of culture area PBS and aspirate to remove residual DTT.
4. Add ~5 µL/mm^2^ 0.25% trypsin (ThermoFisher Scientific) apically and twice the apical volume basolaterally (e.g. for a 6.5 mm insert add 250 µL trypsin apically and 500 µL basolaterally) and incubate cells at 37 °C for ~15 minutes.
5. Harvest trypsinized cells by pipetting up and down gently on the apical surface, then pipetting through a 100 µm filter into prewarmed serum containing medium to neutralize trypsin. Rinse the insert twice using the basolateral trypsin to maximize cell recovery.
6. Centrifuge cells (400 × *g* for 5 minutes).

**Flow cytometry analysis**

1. Resuspend cells in 0.5% (v/v) paraformaldehyde (PFA, ThermoFisher Scientific) diluted in PBS and incubate at room temperature for 8 minutes.
2. Centrifuge cells at 400 × *g* for 5 minutes and wash twice in PBS to remove residual PFA (Note 3).
3. Resuspend in 5% (v/v) normal goat serum (NGS; Jackson ImmunoResearch Inc., West Grove, PA) diluted in PBS containing human TruStain FcX (Biolegend, San Diego, CA) on a rotator for ~15 minutes at room temperature to block non-specific primary antibody binding and Fc receptor binding, respectively.
4. While blocking add appropriate volumes of cell surface primary antibodies to a 96-well plate (Note 4).
5. Add 100,000 blocked cells to each well containing primary antibody and incubate at 4 °C for 30 minutes on a rocker
6. Centrifuge plate(s) and wash cells 2-3× in blocking buffer.
7. After the final wash resuspend cells in blocking buffer containing 0.2% (w/v) saponin (MilliporeSigma) for 15 minutes to simultaneously permeabilize cells and block non-specific binding intracellularly.
8. While permeabilizing, add appropriate volumes of intracellular antibodies to a 96-well plate.
9. Transfer permeabilized cells to plates containing intracellular marker antibodies and incubate at 4 °C for 40 minutes on a rocker.
10. During intracellular staining, prepare compensation controls using UltraComp beads and isotype control antibodies as per the manufacturer’s instructions.
11. Wash cells twice and resuspend in eBioScience Flow Cytometry Staining Buffer (ThermoFisher Scientific).

**SiR Tubulin staining**

1. Prepare 1 µM staining solution by diluting SiR-Tubulin in ALI medium supplemented with 10 µM verapamil (Note 5).
2. Aspirate conditioned ALI medium from cells, replace with SiR-tubulin staining media and incubate for 1 hr at 37 in a humidified atmosphere containing 5% CO2.
3. Continue with ‘Generating HBE single cell suspension’ then ‘Flow cytometry cell sorting’ (immediately below).

**Flow cytometry cell sorting**

1. Resuspend cells at 10^6^ cells/mL in cation-free PBS containing fixable viability dye eFluor 450 (1:2000; eBioscience) in the dark for 10 minutes at 4 °C.
2. Centrifuge cells and wash once in PBS.
3. Centrifuge then block and stain as above (see ‘Flow cytometry analysis’, steps 3-6; note permeabilization and intracellular staining is not performed).

**Data acquisition**

1. Before acquiring the data, set up the flow cytometer. Run CST beads for manual quality control of the instrument. Using fully stained cells, set voltages for all channels. Set up compensation using Ultracomp beads to account for spectral overlap between fluorophores.
2. Isotype controls obtained from the same manufacturer as the primary antibody, conjugated to the corresponding fluorophore as the primary antibody, and used at the same concentration as the primary antibody, should be used to assess nonspecific staining and to set negative gates for flow cytometric analysis. The gating region should be set so that ≤1% of the cells stained with negative controls fall into the positive gate.

**Notes**

1. PBS alone can be used if only unstimulated cells are used, but IL-13 results in tethered MUC5AC which requires reduction for removal (2).
2. The wash can be aspirated or pipetted and stored frozen if investigation of the mucus gel/secreted proteins is warranted.
3. PFA-fixed cells can be frozen at -80°C and used at a later date.
4. FACS tubes can be used alternatively.
5. Verapamil is a broad-spectrum efflux pump inhibitor, reported to greatly improve SiR-tubulin staining .

**References**

1.	Hill DB, Button B. Establishment of Respiratory Air–Liquid Interface Cultures and Their Use in Studying Mucin Production, Secretion, and Function. In: McGuckin MA, Thornton DJ, editors. *Mucins: Methods and Protocols* Totowa, NJ: Humana Press; 2012. p. 245–258.doi:10.1007/978-1-61779-513-8_15.

2.	Bonser LR, Zlock L, Finkbeiner W, Erle DJ. Epithelial tethering of MUC5AC-rich mucus impairs mucociliary transport in asthma. *J Clin Invest* 2016;126:2367–2371.
